## Supplementary Information for "Structure-function analysis of purified proanthocyanidins reveals a role for polymer size in suppressing inflammatory responses"

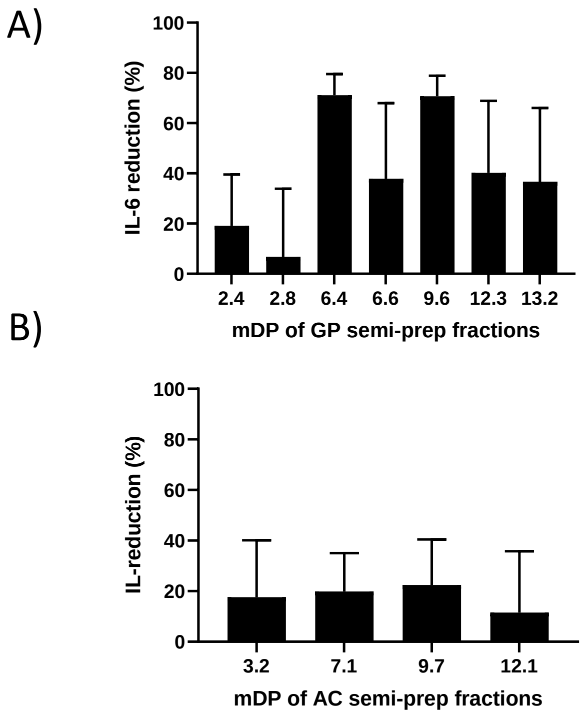


**Supplementary Figure 1 - Inhibition of IL-6 secretion in LPS-activated macrophages stimulated with grape pomace or alpine currant PAC at equimolarity of 7.8 µM.** Experiments were conducted at least twice with triplicate samples.

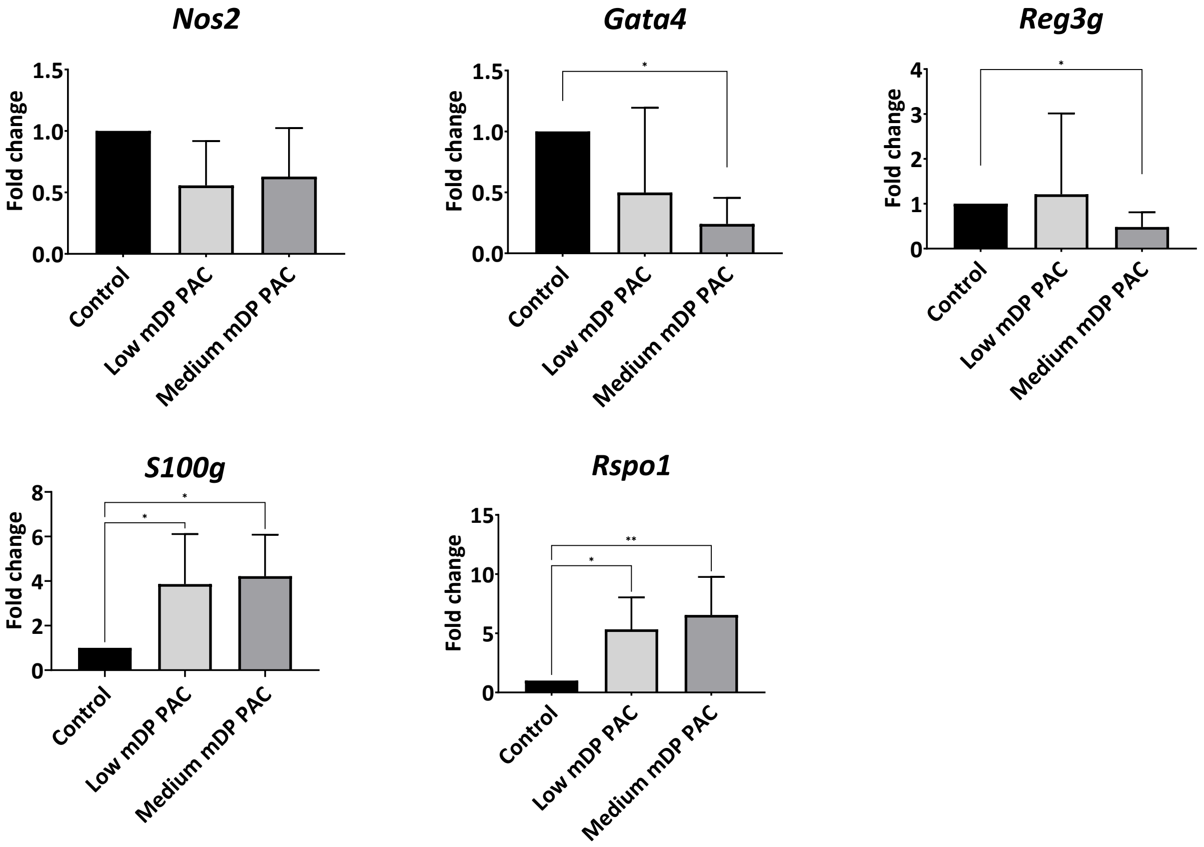


**Supplementary Figure 2 - Regulation of gene expression in mouse ileum tissue by PAC**

qPCR data depicting the regulation of *Nos2*, *Gata4*, *reg3g, S100g* and *Rspo1* in the ileum tissue of mice dosed with either low or medium mDP Sephadex fractions derived from grape pomace (GP). Data is expressed as fold changes relative to mice dosed with water only (*n*= 5 mice per treatment group). (******p* < **0.05, *****p* **< 0.01 by Kruskal-Wallis test).**

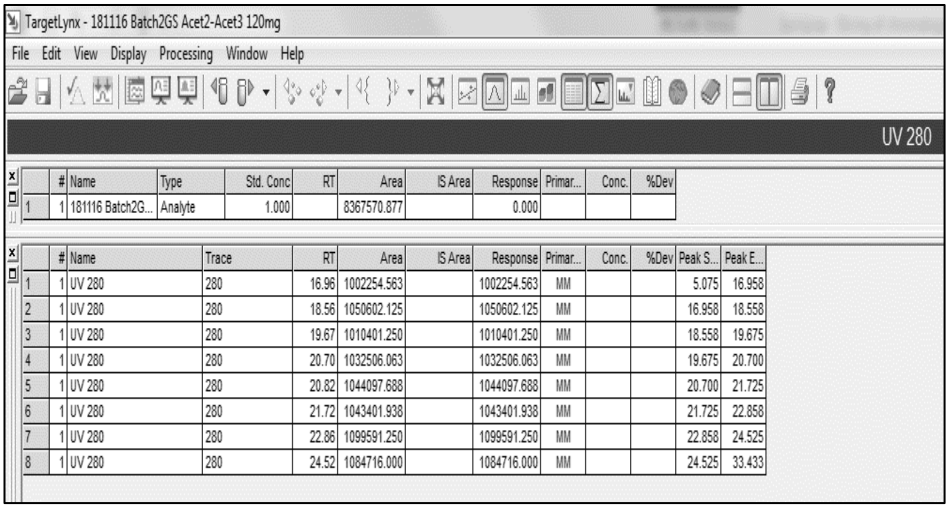

TargetLynx - 181116 Batch2GS Acet2-Acet3 120mg

File Edit View Display Processing Window Help

UV 280

| | # | Name | Type | Std. Conc | RT | Area | IS Area | Response | Primar... | Conc. | %Dev |
|---|---|---|---|---|---|---|---|---|---|---|---|
| 1 | 1 | 181116 Batch2G... | Analyte | 1.000 | | 8367570.877 | | 0.000 | | | |

| | # | Name | Trace | RT | Area | IS Area | Response | Primar... | Conc. | %Dev | Peak S... | Peak E... |
|---|---|---|---|---|---|---|---|---|---|---|---|---|
| 1 | 1 | UV 280 | 280 | 16.96 | 1002254.563 | | 1002254.563 | MM | | | 5.075 | 16.958 |
| 2 | 1 | UV 280 | 280 | 18.56 | 1050602.125 | | 1050602.125 | MM | | | 16.958 | 18.558 |
| 3 | 1 | UV 280 | 280 | 19.67 | 1010401.250 | | 1010401.250 | MM | | | 18.558 | 19.675 |
| 4 | 1 | UV 280 | 280 | 20.70 | 1032506.063 | | 1032506.063 | MM | | | 19.675 | 20.700 |
| 5 | 1 | UV 280 | 280 | 20.82 | 1044097.688 | | 1044097.688 | MM | | | 20.700 | 21.725 |
| 6 | 1 | UV 280 | 280 | 21.72 | 1043401.938 | | 1043401.938 | MM | | | 21.725 | 22.858 |
| 7 | 1 | UV 280 | 280 | 22.86 | 1099591.250 | | 1099591.250 | MM | | | 22.858 | 24.525 |
| 8 | 1 | UV 280 | 280 | 24.52 | 1084716.000 | | 1084716.000 | MM | | | 24.525 | 33.433 |

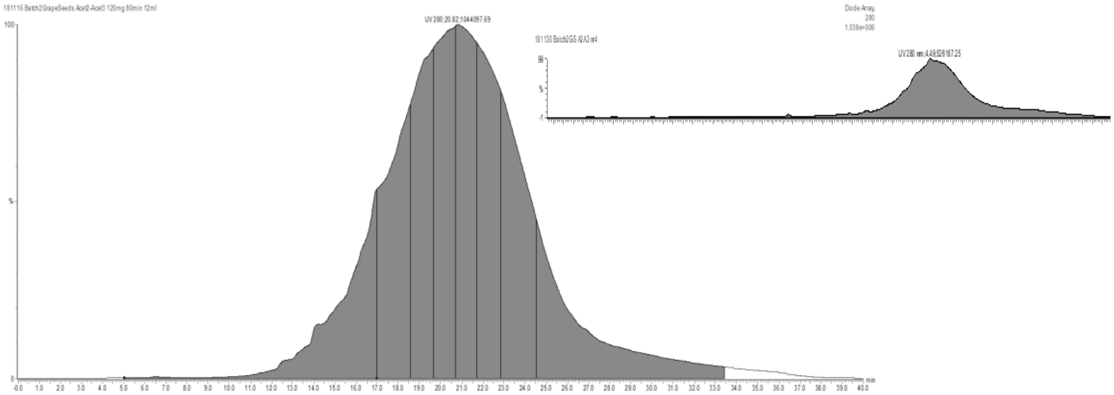

**Supplementary Figure 3 - Integration of Chromatograms**

MassLynx was used in order to determine the Area under the curve (AUC) within the time-span of 5–33 minutes. The Peak Response Area value was divided into 8 equally sized “slices”, corresponding to the 8 areas of the chromatogram. The 168 tubes resulting from the semi-preparative liquid-chromatography were then pooled accordingly into 8 highly purified PAC samples, which were individually analyzed by UPLC-MS/MS.

### Supplementary Table 1 - List of material used for the extraction and purification of proanthocyanidins

| | Material | References |
|---|---|---|
| Samples | Alpine currant (AC) Macerated in 80 % analytical acetone (09-08-2018) | This study |
| | Grape pomace (GP) Macerated in 80 % analytical acetone (14-09-2018) | Nor-Feed A/S (Hvidovre, Denmark) |
| Solvents | Acetyl acetate | VWR International S.A.S, France |
| | Analytical acetone | VWR International S.A.S, France |
| | Butanol | VWR International S.A.S, France |
| | Diethyl ether | VWR International S.A.S, France |
| | Ethyl acetate | VWR International S.A.S, France |
| | Formic acid 99-100% | WR International, EC |
| | Formic acid, LC-MS grade | Sigma Aldrich, USA |
| | Methanol, analytical grade | VWR International S.A.S, France |
| | Acetonitrile, LC-MS grade | VWR International S.A.S. (USA) |
| Other | MilliQ water Purified with Millipore Synergy UV system | Merck KGaA, Darmstadt, Germany |
| | Sephadex LH-20 | GE Healthcare |

**Supplementary Table 2 - Extraction and purification of proanthocyanidins**

Overview of the extraction and purification steps used to isolate purified PAC from grape pomace (GP) and alpine currant (AC) by Sephadex LH-20 fractionation followed by semi-preparative liquid chromatography.

| | Alpine currant | Grape pomace |
|---|---|---|
| 1. Extraction by filtration | Macerated in 80 % analytical acetone (09-08-2018) | Macerated in 80 % analytical acetone (14-08-2018) |
| | 5 extractions through Büchner funnel | |
| | Evaporation of acetone from samples in fume hood | |
| | UPLC analysis | |
| | Pooling of extractions 1-5 | Pooling of extractions 1-5 |
| | Liquid-liquid extraction with ethyl acetate and butanol | - |
| | - | - |
| 2. Sephadex fractionation | Sephadex fractionation<br>6 water fractions<br>5 methanol fractions<br>6 acetone fraction | Sephadex fractionation<br>6 water fractions<br>5 methanol fractions<br>6 acetone fraction |
| | O/N evaporation of acetone fractions<br>Rotary evaporation of methanol fractions | |
| | Freeze-drying and weighing of samples | |
| | Pooling of samples based on similarity of chromatogram → 6 alpine currant sephadex samples | Pooling of samples based on similarity of chromatogram → 8 grape pomace sephadex samples |
| 3. Semi-preparative liquid chromatography | Injection of ~120 mg of each sephadex fractions | |
| | Collection of samples in 168 Eppendorf tubes | |
| | Pooling of each semi-prep run into 8 equal sub-fractions → 48 alpine currant semi-prep fractions | Pooling of each semi-prep run into 8 equal sub-fractions → 64 grape pomace semi-prep fractions |
| | Evaporation of solvents by rotavap, rotary evaporation or centrifugal concentration<br>and freeze-drying | |
| | A total of 112 semi-prep samples containing highly purified PAC were generated with sample<br>weights ranging between 2-17 mg | |

### Supplementary Table 2 - Primer sequences used in experiments

| Primer name | Primer sequence (5'-> 3') |
|---|---|
| **Used for *in-vitro* studies** | |
| *Tlr4* forward primer | *ACTGGCCTTTCAGGAACTTT* |
| *Tlr4* reverse primer | *ACATCCTAGGGCTGTCTTTCTT* |
| *Atp6v0d2* forward primer | *GGGCCAGTGTTCAGTTGCTA* |
| *Atp6v0d2* reverse primer | *TCCTGCTGAGTTAGGAGGCT* |
| *Rab7b* forward primer | *GGAAGTGGCCTCTCACCAAA* |
| *Rab7b* reverse primer | *CCTCACACAGGTGGGAGTTC* |
| *Gapdh* forward primer | *TATGTCGTGGAGTCTACTGGT* |
| *Gapdh* reverse primer | *GAGTTGTCATATTTCTCGTGG* |
| **Used for *in-vivo* studies** | |
| *Nos2* forward primer | *GGTGAAGGGACTGAGCTGTT* |
| *Nos2* reverse primer | *TGCACTTCTGCTCCAAATCCA* |
| *Gata4* forward primer | *TTCTGGGAAACTGGAGCTGG* |
| *Gata4* reverse primer | *TGCTTTCTGCCTGCTACACA* |
| *Reg3g* forward primer | *CACCATCCTAGGGATCTGCAA* |
| *Reg3g* reverse primer | *ATGGGGCATCTTTCTTGGCA* |
| *S100g* forward primer | *GGAGCTGGATAAGAATGGCGA* |
| *S100g* reverse primer | *AGAGCGTGCGTTCAATCAGT* |
| *Rspo1* forward primer | *TGTACTTACACAAGGGCCGC* |
| *Rspo1* reverser primer | *GGGACCACTCGCTCATTTCA* |
| *Gapdh* forward primer | *TATGTCGTGGAGTCTACTGGT* |
| *Gapdh* reverse primer | *GAGTTGTCATATTTCTCGTGG* |
